# Monoclonal antibody to Filarial serpin Wb123 impedes the urokinase plasminogen activator receptor mediated Alternative activation of macrophages

**DOI:** 10.1101/2025.02.05.636671

**Authors:** Prince Upadhyay, Akshay Munjal, Abir Mondal, Gagandeep Singh, Soumyadeep Mukherjee, Mahesh C Kaushik, Puneet K. Gupta, Soumya Pati, Shailja Singh

## Abstract

Macrophages play a multifaceted role in mounting an immune response against pathogens through phagocytosis, antigen presentation and cytokine production. Activated macrophages are critical for the initiation, maintenance, and resolution of inflammation. They can either be classically activated (Th1 response), which promotes inflammation or alternatively activated (Th2 response), which resolves inflammation. The alternate activation of macrophages, along with diminished T cell response, is thought to be an important immune evasion strategy employed by the filarial parasites. However, the mechanism behind filaria-induced alternate activation remains unclear. In this study, by using in-silico approach we identified filarial serpins which are highly expressed in the infective L3 larval stages. We characterized the filarial serine protease inhibitor, Wb123 and established its role in alternate activation. We observed that Wb123 impairs nitric oxide (NO) and reactive oxygen species (ROS) expression, which plays a crucial role in mounting a strong immune response. Surprisingly, we also observed elevated IL6 expression and phosphorylation of STAT3, indicating IL-6-dependent alternate activation. Our findings indicate that the urokinase plasminogen activator receptor (uPAR) is essential for Wb123-induced alternate activation. Furthermore, we demonstrated that the monoclonal antibody MAbG8 impedes the Wb123-induced alternate activation, as shown by reduced CD163 expression along with increased ROS and uPA expression in response to lipopolysaccharide (LPS) and interferon-gamma (IFN-γ). Overall, our study identifies the filarial serpin Wb123 and highlights its potent role in alternative activation. This interaction reveals, for the first time a mechanism exploited by filarial parasites to evade a strong pro-inflammatory immune response.

## 1. Introduction

Lymphatic filariasis is primarily caused by worms such as *Wuchereria bancrofti*, *Brugia malayi* and *Brugia timori*, while filarial worms like *Onchocerca volvulus* and *Loa loa* causes subcutaneous filariasis.^1^ Lymphatic filariasis is a neglected tropical disease endemic to 39 countries, affecting approximately 36 million people with chronic disease manifestations and putting an additional 657 million at risk of infection.^2^ These worms can live for about 6 to 8 years, producing millions of microfilariae that migrate to various organs, including the brain.^3,4^ Microfilaria elicit a modified type 2 immune response characterized by the production of cytokines like IL-4, IL-10 and IL-13 along with the expansion of regulatory T cells.^5,6^ Further studies on infective stage (L3 larvae) have shown reduced CD4+ T cell activation due to decreased ability of Langerhans’ cells (resident macrophages), indicating altered antigen-presenting function.^7^ Macrophages are an important class of antigen-presenting cells that play a crucial role in providing protection against pathogens by producing nitric oxide and other mediators. They alter their phenotype and function in response to microenvironment stimuli, acquiring either a classically or alternatively activated phenotype.^8^ Classically activated macrophages are essential components of innate immune response that are detrimental to filarial parasites. These macrophages express molecular markers like CD86, CD68 and Toll like receptor (TLR4) while secreting various pro-inflammatory cytokines like IL-1β, IL-12, ROS and NO.^9^ A different class of macrophages are induced in response to filarial infection and filaria-derived antigens, known as alternatively activated macrophages characterized by increased expression of IL-10 and arginase 1 and along with markers like CD163.^10^ Studies have highlighted that filarial patients have monocyte studded with filarial antigens throughout the course of infection, suggesting the influence of secreted proteins commonly referred to as excretory-secretory products (ESP) and have been demonstrated to modulate lymphatic function and significantly alter monocyte activity.^11,12^ A comparative analysis of ESP obtained from microfilariae, male and female worms revealed that homologues of serine protease inhibitors (serpins) were highly abundant in microfilaria ESP.^13,14^ Serpins are a superfamily of protease inhibitors found ubiquitously in organisms ranging from viruses to vertebrates. They possess a highly conserved secondary structure averaging 350-400 amino acids with a distinct reactive center loop (RCL), an exposed protrusion of 20-25 amino acids. RCL of serpins contains recognition sites (P1-P1’) that determine the specificity of each serpin for serine proteases.^15,16^ Recently, a study identified a serpin from *Trichinella spiralis* and highlighted its role in the alternate activation of macrophages.^17^ This discovery raises questions about the functional roles and molecular mechanisms of filarial serpins. Consequently, we investigated the potential role of filarial serpins as modulators of the host immune response.

## 2. Methods

### Culturing of Human brain macrophage

Human brain macrophage cells HMC3 (Human microglia clone 3) was ordered from ATCC (CRL-3304). Cells were cultured in EMEM supplemented with 10% FBS, 1 mM Sodium Pyruvate, 1x NEAA, and 0.1% Penicillin-streptomycin. The culture was maintained in a 25cm^2^ Corning flask under humid conditions at 37 ℃ with 5% CO_2_. Sub-confluent HMC3 cells were passaged every second day using 0.5% Trypsin-EDTA. To induce classical activation of microglia, LPS (1ug/ml) was used along with IFN-γ (100ng/ml). All experiments were performed within 4 to 15 passages.

### Immunofluorescence of microfilaria

Microfilaria positive (MF+) filarial blood samples were diluted with 1xPBS. Cells were pelleted and fixed using 4% paraformaldehyde for 20 minutes at room temperature, then washed with 1xPBS. Blocking was performed with 4% BSA for 2 hours at room temperature. Primary monoclonal and polyclonal antibodies against Wb123 were incubated at 4℃ for 2 hours, followed by washing with a blocking buffer. An Alexa Fluor-488 (Biorad, #5196-2404) conjugated secondary antibody was added and incubated for 1hour before washing with 1xPBS. Blood smears were mounted using Vectashield antifade DAPI (H-1200-10) mounting media, and images were captured using a Nikon Ti2 confocal microscope.

### Isolation of PBMCs from filarial patients

After initially screening multiple individuals from the Govindpur region of Dhanbad district in Jharkhand. A field-based immunochromatographic card test for circulating filarial antigen was performed using whole blood collected from five individuals between 9 p.m. and midnight. Microscopic examination of calibrated thick smears resulted in the identification of two microfilaria-positive (MF+) and three microfilaria-negative (MF-) samples. Peripheral blood mononuclear cells (PBMC) from the blood of MF+ filarial patients and endemic normal donors were isolated using Ficoll density centrifugation (Hi-Sep LSM 1077). Trizol reagent was added to isolate RNA and cDNA was subsequently used to perform qPCR. All study participants provided informed consent and were examined as a part of protocols approved by review board of TCI foundation (NBS_Godda-2023-24-32859).

### Cloning and purification of rWb123 and antibody production

The Coding sequence (CDS) of the Wb123 gene was codon-optimized, synthesized de novo, and subcloned into NcoI and XhoI digested pET-28a (+) vector and transformed in E. coli Rosetta (DE3) cells. The expression of recombinant protein was induced using 1mM IPTG at 18℃ for 16 hours. The cell biomass was pelleted at 8000 g for 10 minutes, followed by lysis and sonication in resuspension buffer (30mM Tris-Cl, 0.5mM EDTA pH 7.5) until the viscous fluid became clear. Inclusion bodies were pelleted and washed repeatedly with buffer (50mM Tris-Cl and 100mM NaCl, 0.5% triton-100 and 0.1% sodium azide) followed by a final wash with buffer containing 50mM Tris (pH 8.0) and 500mM NaCl. The pellet was dissolved in a buffer containing 6M GuHCl, 10mM Tris (pH 8.0), 300 mM NaCl at room temperature for 16 hours. Following incubation, cells were centrifuged (Thermo-scientific, sorvall, LYNX-4000) at 13,000 rpm for 30 minutes at 4℃. Ni-NTA beads were incubated with the supernatant on a rotor shaker overnight at 4℃. The resins were packed into a column and washed with ten volumes of 8 M urea, 20 mM Tris-Cl and 500mM NaCl (pH 8.0) containing 20mM imidazole. Protein elution fractions were collected at imidazole concentrations of 50mM, 150mM, 250mM and dialyzed in 20 mM Tris-HCl with NaCl at 150mM (pH 7.4), 1mM PMSF. The dialyzed protein concentration was determined by BCA assay and then run on a 15% SDS-PAGE.

### Immunocytochemistry assay

To analyze the intracellular expression of proteins after treatment with rWb123 and LPS/IFN-γ in different combinations (Table 1), cells were seeded on the coverslips at a density of 1.5 x 104 in 24 well plate (NEST, tissue culture). After treatment, cells were fixed by 4% paraformaldehyde followed by permeabilization using 0.10% Triton X-100 and blocking with 2% BSA for 1 h prepared in 1x PBS. Rabbit raised primary antibody dilutions were incubated overnight at 4℃. Alexa Fluor-488 conjugated anti-rabbit were used as secondary antibody. Vectashield antifade DAPI (H-1200-10) mounting media was used to mount the coverslips on the shield.

**Table 1:** Experimental strategies for marker expression analysis.

| **S. No.** | **Treatment**<br>**(5 Hours)** |
| --- | --- |
| 1. | Control<br>(Media) |
| 2. | Condition 1<br>(rWb123) |
| 3. | Condition 2<br>LPS and IFN-γ (LPS-I), 1ug/ml-100ng/ml) |
| 4. | Condition 3<br>(LPS-I+rWb123, 250ng) |

### In silico screening-Molecular docking and simulation analysis

The proteome of filarial worms was compiled using FASTA protein files of *Wuchereria bancrofti* (BioProject-PRJEB536), *Brugia malayi* (BioProject-PRJNA10729), *Brugia timori* (BioProject-PRJEB4663) and *Onchocerca volvulus* (BioProject-PRJEB513), downloaded from WormBase Parasite database. The raw HMM sequence for serine protease inhibitors was obtained from the Pfam webserver (PF00079) and a standalone BLAST search was conducted against the compiled database. Multiple sequence alignment was performed using Clustal omega. The predicted structure of Wb123 was retrieved from AlphaFold Protein Structure Database and validated through a Ramachandran plot (Saves-v6.1) and ProSA-web server. The crystal structure of human uPA (PDB ID: 4DVA) at 1.94Å resolution was retrieved from the Protein DataBank (PDB). Molecular docking of Wb123 and uPA was performed by using HDOCK server. Protein-protein complex interactions were evaluated with the PDBsum and LigPlus software. The uPA-Wb123 complex stability was analyzed using GROMACS 5.1.5 through the LiGRO tool. The system was prepared in a TIP3P water model with 0.15 M NaCl in a cubic box (4 nm periodic image distance) using the Amber99sb force field. System equilibration involved NVT and NPT ensemble runs (1000ps each) using Berendsen thermostat and Parrinello-Rahman barostat. A 150 ns production MD was performed at 310.15 K and 1.01 bars. The simulation employed PME for electrostatics, LINCS for hydrogen bonds, and a 2fs time step with 1ps frame storage. Analysis utilized PyMOL and VMD for visualization, standard GROMACS tools for stability metrics, and gmx_MMPBSA tool for binding energy calculations from the final 150 frames.

### Antibody-dependent blocking of urokinase plasminogen activator receptor

To evaluate the effect of uPAR blocking on rWb123 function, we pre-incubated cells in one well with uPAR antibody (DF12495) and one well with LRP1 (DF2935) for 2 hours. rWb123 was incubated for 5 hours. After incubation, cells were lysed using RIPA buffer and run on 12% SDS-PAGE to evaluate CD163 expression using immunofluorescence and western blot assay.

### Western Blotting

To investigate pro-inflammatory and anti-inflammatory markers expression in conditions described in Table 1. Cells were lysed using RIPA buffer containing 1x Protease inhibitor cocktail (Roche). Protein estimation was performed using the BCA assay (Sigma, #QPBCA-1KT). Equal amount of protein from each sample were separated on 12% SDS-PAGE (Biorad) and was transferred to the PVDF membrane using Trans-blotter (Biorad). The blots were evaluated for protein expression using primary antibodies shown in table 4 followed by 1h anti-rabbit HRP conjugated secondary antibody (1:5000) incubation. GAPDH was used as loading control. Blots were developed in iBright (Invitrogen) using ECL substrate (Biorad #1705061).

### Enzyme-Linked immune sorbent assay (ELISA) to evaluate MAbG8 interaction with Wb123

To assess the interaction of Wb123 with MAbG8, a 96-well ELISA plate was coated with purified Wb123 (bait; 100 ng of each) in PBS at RT for 5 h, blocked with 5% BSA in PBS overnight at 4°C. Further, increasing concentration of MAbG8 was added and incubated for 2 h at RT. After washing with PBS, secondary anti-mice HRP conjugated antibody was incubated at RT for 2 h. Detection reagent (TMB; HIMEDIA) was added, after washing with PBS, the HRP reaction was stopped by adding 3M HCl. Absorbance was measured at 450 nm using a microplate reader and graph was plotted using GraphPad PRISM software.

### Antibody neutralization experiments

To evaluate the effect of anti-Wb123 monoclonal antibody (MabG8) on rWb123 induced alternate expression we performed antibody neutralization assay. Here anti-Wb123 antibody (1:5) was incubated with 1ug rWb123 and alone rWb123 was also kept at 4℃ overnight. Next day, Wb123-MAbG8 complex and rWb123 were added to the cells in EMEM media for 5 hours. Then, cells were either fixed using 4% paraformaldehyde for immunofluorescence study or lysed for western blot analysis as described earlier. Similarly, to evaluate the effect on ROS production, cells were treated with LPS/IFN-γ for 3 hours before addition of Wb123-MAbG8 complex and rWb123 in EMEM media. 5µM carboxy-H2DCFDA (Invitrogen catalog #136007) was added to the cells in Opti-MEM media for 20 minutes and visualized using confocal microscopy. We also evaluated the uPA expression using anti-uPA antibody (1:200).

### Study the ROS by flow cytometer

For ROS expression analysis cells were given treatment as described (Table-2). After treatment cells were trypsinized and centrifuged (Gyrogen-1580R) at 850rpm for 5 mins followed by wash using 1x PBS. Afterwards cells were suspended in 200ul of 1x PBS and 100nM carboxy-H2DCFDA (Invitrogen, #136007) was added to the 1.5 ml microcentrifuge tubes. We used 20µM H_2_O_2_ (10 min) as a positive control and no-dye control to nullify the background noise in our experiment. We performed flow cytometry in Beckman Cytoflex flow cytometer and 10000 events were captured.

**Table 2:**
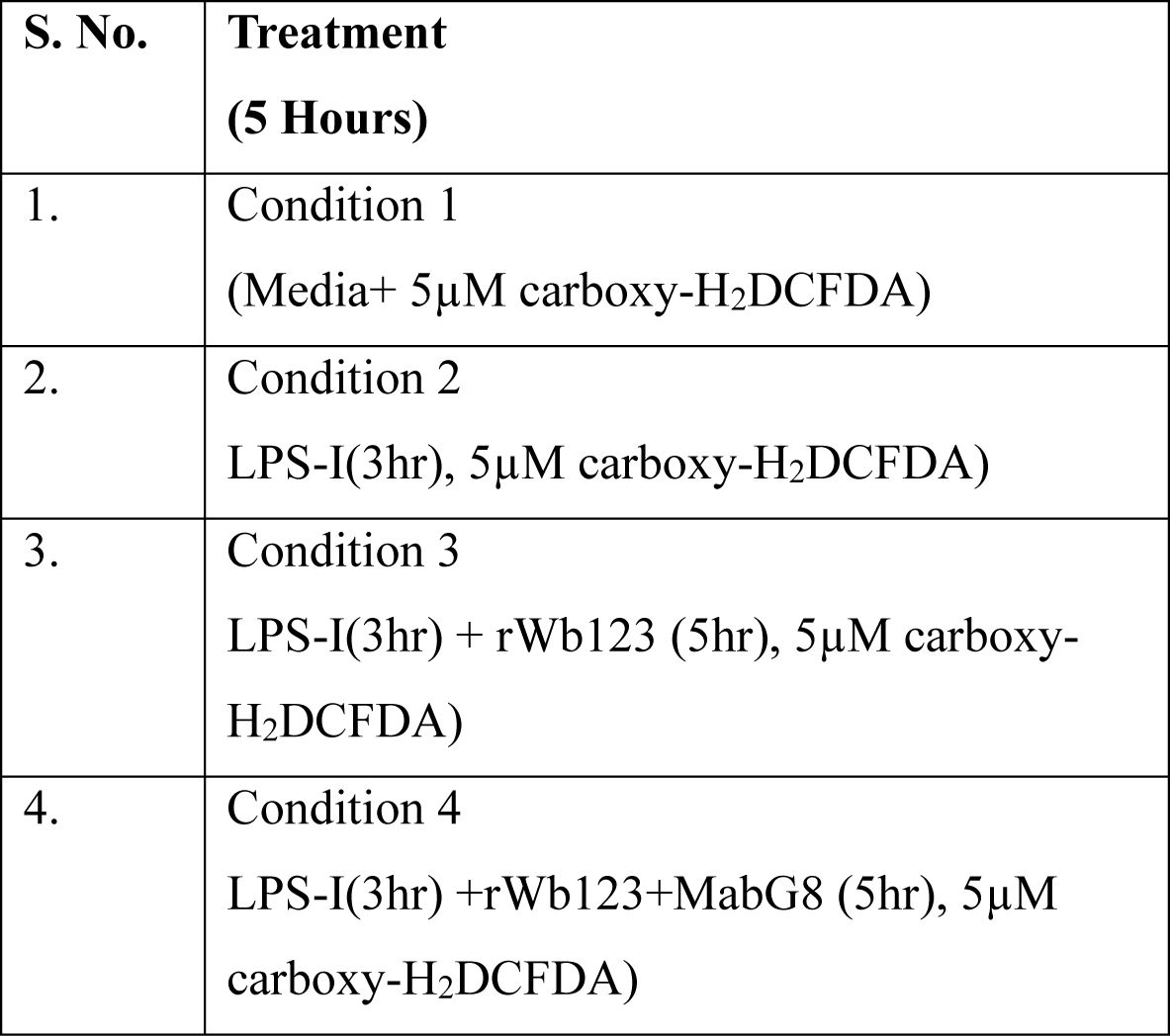
Flowcytometry strategies for ROS expression.

| **S. No.** | **Treatment**<br>**(5 Hours)** |
| --- | --- |
| 1. | Condition 1<br>(Media+ 5μM carboxy-H<sub>2</sub>DCFDA) |
| 2. | Condition 2<br>LPS-I(3hr), 5μM carboxy-H<sub>2</sub>DCFDA) |
| 3. | Condition 3<br>LPS-I(3hr) + rWb123 (5hr), 5μM carboxy-H<sub>2</sub>DCFDA) |
| 4. | Condition 4<br>LPS-I(3hr) +rWb123+MabG8 (5hr), 5μM carboxy-H<sub>2</sub>DCFDA) |

Similarly, for studying CD163 expression, we incubated cells with anti-CD163 primary antibody (1:200) for 2 h at RT followed by Alexa Fluor-488 (Bio-Rad, #5196-2404) conjugated anti-rabbit secondary antibody incubation for 1h. Flow cytometry was performed and the stacked histogram was plotted using Floreada.io.

### Q-PCR-based gene expression study

Microglia cells after 5 hours treatment in various experimental conditions were trypsinized and pellet down to isolate RNA. TRIZOL reagent (Thermofisher, #15596206) was used and the RNA concentration was measured using the Nanodrop 2000 instrument (Thermo-Scientific). The RNA was used to make complementary DNA (cDNA) using cDNA reverse transcription kit (Applied Biosystems, #4368814). Powerup SYBR green master mix (Applied Biosystems # A25742) and gene-specific primers were used to perform quantitative PCR in AB-Step OnePlus instrument. Fold change (2-ΔΔCt) calculations were performed in Microsoft excel, normalized with the housekeeping gene 18S. The bar plot was plotted using GraphPad Prism 9 software. The qPCR primers are mentioned. (Table 3)

**Table 3:**
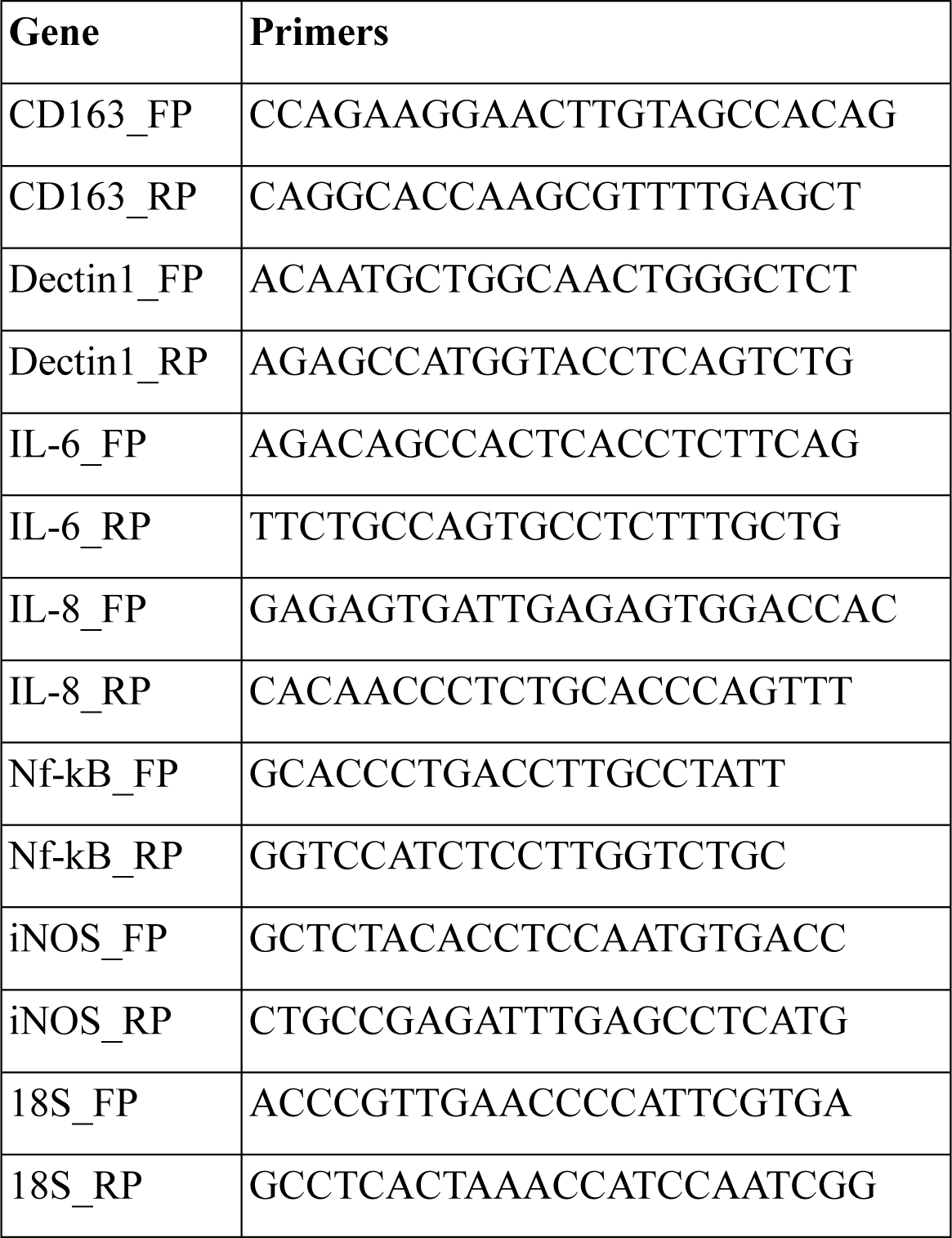
Primers details.

**Table 4:** Antibody details.

| **Antibodies** | **Identifier** |
| --- | --- |
| CD163 | Cat #DF8235 |
| CD86 | Cat #DF6332 |
| STAT3 | Cat #AF6294 |
| PSTAT3 (Tyr705) | Cat #AF3293 |
| PLAU | Cat #DF6904 |
| ARG1 | Cat #DF6657 |
| PLAUR | Cat #DF12495 |
| LAMIN | Cat #AF6056 |
| LRP1 | Cat #DF2935 |
| GAPDH | Cat #AF0911 |
| Anti-His | Cat # pAb-T0051 |
| Anti-rabbit HRP 2° | Cat #31460 |
| Anti-mouse HRP 2° | Cat #31430 |
| Alexa Fluor-488 | Cat #5196-2404 |

### uPA inhibition assay

Inhibition of uPA activity was tested by performing direct fluorescent assay. Here, serial dilutions of Wb123 were added to the wells of 96-well plate (black) on ice containing recombinant uPA from human kidney cells (Cat# SRP6273, Sigma-Aldrich) and urokinase fluorogenic substrate III (Z-Gly-Gly-Arg-AMC, Cat# 672159, Sigma-Aldrich) to make a final volume of 200 μL/well, final enzyme concentration 2 nM, and final substrate concentration 250 μM in assay buffer(20 mM HEPES pH 7.4, 100 mM NaCl, 0.5 mM EDTA, 0.01% v/v Tween-20). Reaction was monitored immediately following addition of the enzyme using a microplate fluorescence reader (BioTek, Synergy H1) using excitation wavelength (365-380 nm) and emission wavelength (430-460 nm). Changes in fluorescence were observed by incubating after every 2-minute period at 37 °C occurring over 2 h. Significant inhibition of the uPA activity by the different Wb123 concentrations compared to uPA alone. Recombinant PAI1(Cat# A8111, Sigma-Aldrich) was used in 1:1 ratio to uPA as positive control. Graphed using GraphPad PRISM v8.0 software.

## 3. Results

### In-silico screening identifies filarial serpins highly expressed by infective L3 larva and microfilaria

A previous study reported that serpins are part of excretory and secretory products of filarial worms and are highly expressed by microfilariae compared to adult worms.^14^ In the present study, we performed virtual screening using a filarial proteomic database containing 91341 transcripts. To identify extracellular filarial serpins, we began by annotating a pool of proteins for the serpin signature motif (PS00284, PF00079). A total of 66 “high confidence” serpin proteins were retrieved and further annotated for subcellular localization. we subjected all predicted 40 extracellular serpins to conserved domain analysis to screen for serpins with RCL and resulting in the identification of 15 filarial serpins with a conserved RCL. (Fig. 1A) Interestingly, analysis of *Brugia malayi* RNA-seq expression data from WormBase revealed that the expression of serpins was significantly elevated in the infective L3 larva and the microfilaria stage. (Supplementary Fig.1, A) Additionally, we performed multiple sequence alignment of RCL to annotate P1:P1’ residues (scissile bonds) known for their specificity for protease targets. This analysis revealed that 10 out of the 15 serpins have Arginine as a P1 residue, and 5 of these serpins possess both Arginine and Methionine as a P1:P1” residues. (Fig. 1B) This suggests that their target serine protease could be plasminogen activator, which are known to be crucial for activating the complement system and mounting an inflammatory response.^18^

**Figure 1:**
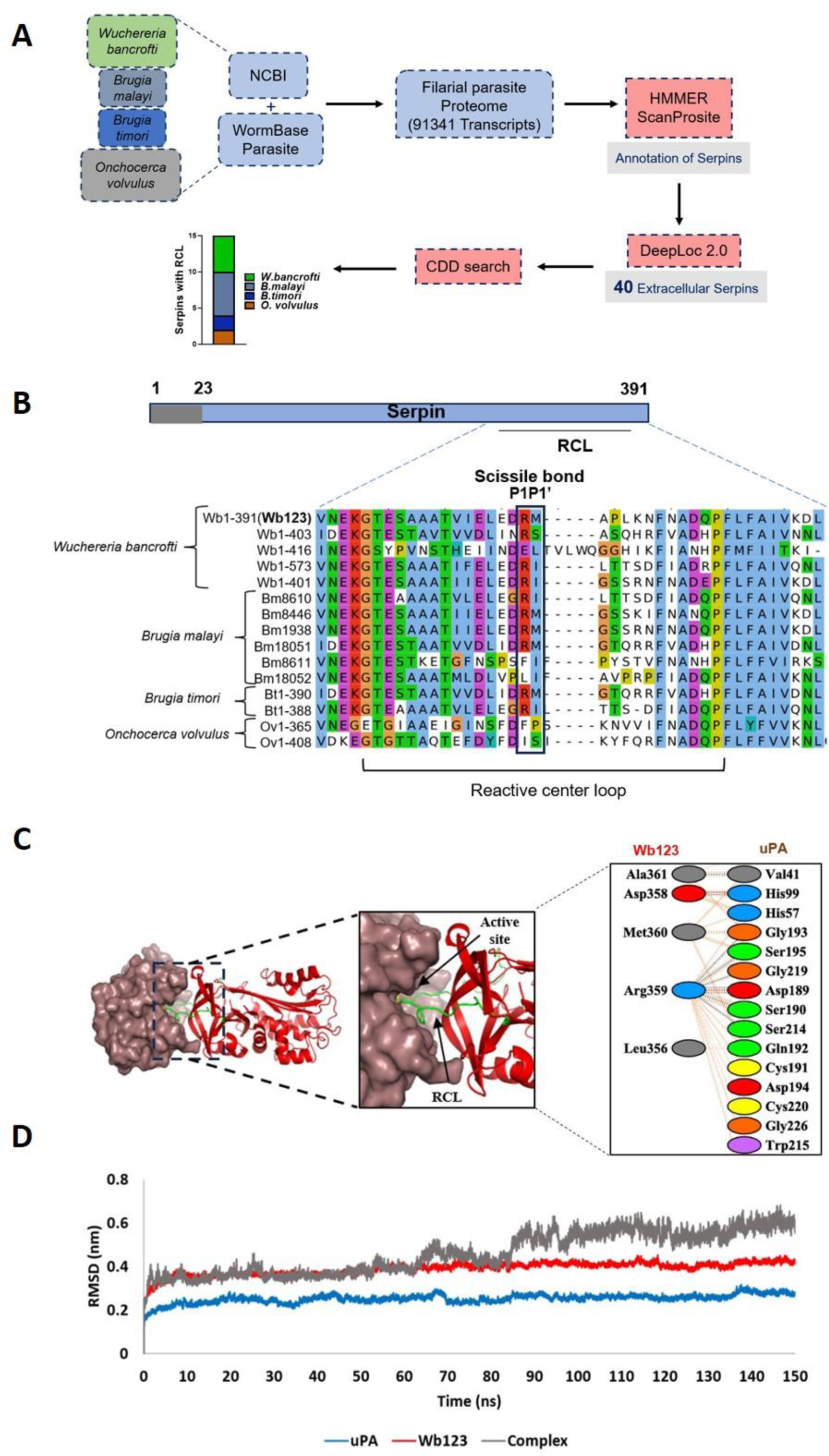
In-silico study identifies filarial serpins and target serine protease. A) Schematic representation of the pipeline devised to screen filarial serpins; B) Multiple sequence alignment of RCL for Wb123 and other identified filarial serpins. P1-P1’ sites are shown in box, The first 2 letters in each serpins represents abbreviations of the scientific names of given species: Wb, *W. bancrofti*; Bm, *B. malayi*; Bt, *B. timori;* Ov; *O. volvulus;* C) *Molecular docking and simulation study.* Representation of molecular docking showing the interaction of filarial serpin Wb123 RCL residues (Arg359, Met360) with catalytic site (His57, Asp189, Ser195) of uPA; D) RMSD graph of Wb123, uPA and Wb123-uPA complex representing 150ns simulation.

### Serine protease plasminogen activator identified as interacting partner of filarial serpin Wb123

Considering that *Wuchereria bancrofti* accounts for more than 90% of filarial cases, we selected the serpin Wb1-391 for further characterization and named it Wb123. The full-length sequence of Wb123 has 391 amino acids, with a theoretical molecular weight of 44kDa and an isoelectric point (PI) of 8.43. Further sequence analysis revealed that wb123 contains a serpin domain (cd00172) with a conserved RCL at positions 345 to 371 aa and an exposed putative nuclear localization signal (PKRRFG) at positions 254 to 259 aa. The modelled structure of Wb123 was validated using the SAVESv6.1 and ProSA web server, yielding an ERRAT score of 93.39. Ramachandran plot indicated that 92.2% (319aa) of residues were in the most favored regions, 6.9% (24aa) in the additionally allowed regions, 0.6% (2aa) in generously allowed regions and 0.3% (1aa) in the disallowed region, with z score of −8.4. Furthermore, the modelled structure of Wb123 was superimposed on resolved structures of PAI-1, Alpha-1-Antitrypsin, Glia-derived nexin and kallistatin showing RMSD values of 1.530, 1.687, 1.611 and 1.845 respectively, confirming structural similarities with human serpins. (Supplementary table1,2)

To investigate serine protease target of filarial serpin Wb123 we performed molecular docking with human serine proteases such as cathepsin G (CTG), plasmin (PL), elastase (EL) and urokinase plasminogen activator (uPA). Docking scores suggested that Wb123 exhibited the highest binding energy of −387.17 for uPA compared to other serine proteases, CTG (−365.01), PL (−295.62) and EL (−308.57). Notably, the RCL was only involved in Wb123-uPA binding. Interaction analysis between Wb123 and uPA revealed that amino acid residues arginine-339 and methionine-340 (p1-p1’) formed a stable complex with the catalytic triad (His57-Ser195-Asp189) of uPA, stabilized by four salt bridges and fourteen hydrogen bonds.^19^ (Fig. 1C) To confirm the structural stability and interaction dynamics of uPA-Wb123 complex we performed molecular dynamics simulation. The RMSD analysis of 150 ns MD simulation revealed distinct stability patterns. uPA showed highest stability (≈0.25 nm RMSD after 10 ns), while Wb123 exhibited slightly higher values (≈0.4 nm), indicating greater flexibility. The uPA-Wb123 complex demonstrated the highest RMSD (0.5-0.6 nm after 90 ns), reflecting combined protein movements while maintaining a stable binding configuration. (Fig. 1D) MMGBSA calculations yielded a binding free energy of −85.38 ± 11.92 kcal/mol, indicating strong complex stability. The RMSD patterns and binding energy collectively confirmed a thermodynamically stable protein-protein interaction throughout the simulation period.

### Recombinant Wb123 induces alternate activation of macrophages

Patient with filarial infections exhibit a compromised immune response characterized by a diminished inflammatory response and an increased frequency of T regulatory cells. ^10^ To understand the activation status of macrophages in filarial patients, we analyzed publicly available gene expression database. Our findings revealed that filarial patients exhibited significantly higher levels of alternative activation markers, including CD163, ARG-1, IL-4 and TGF-β while showing reduced expression of classical activation markers such as CD14 and CD86. Notably, these expression patterns were reversed following treatment, suggesting a potential restoration of immune function. (Supplementary Fig.1B) Additionally, our qPCR results from PBMCs of microfilaria-positive (MF+) patient samples (FP-1 and FP-2) demonstrated increased CD163 and IL-6 expression compared with endemic normal control (EN-1 and EN-2), further solidifying the alternate activation status. (Fig. 2A) Interestingly, a study by Kubala et. al demonstrated that human serpin E1 promotes the alternative activation of macrophages, as evidenced by increased CD163 expression.^20^

**Figure 2:**
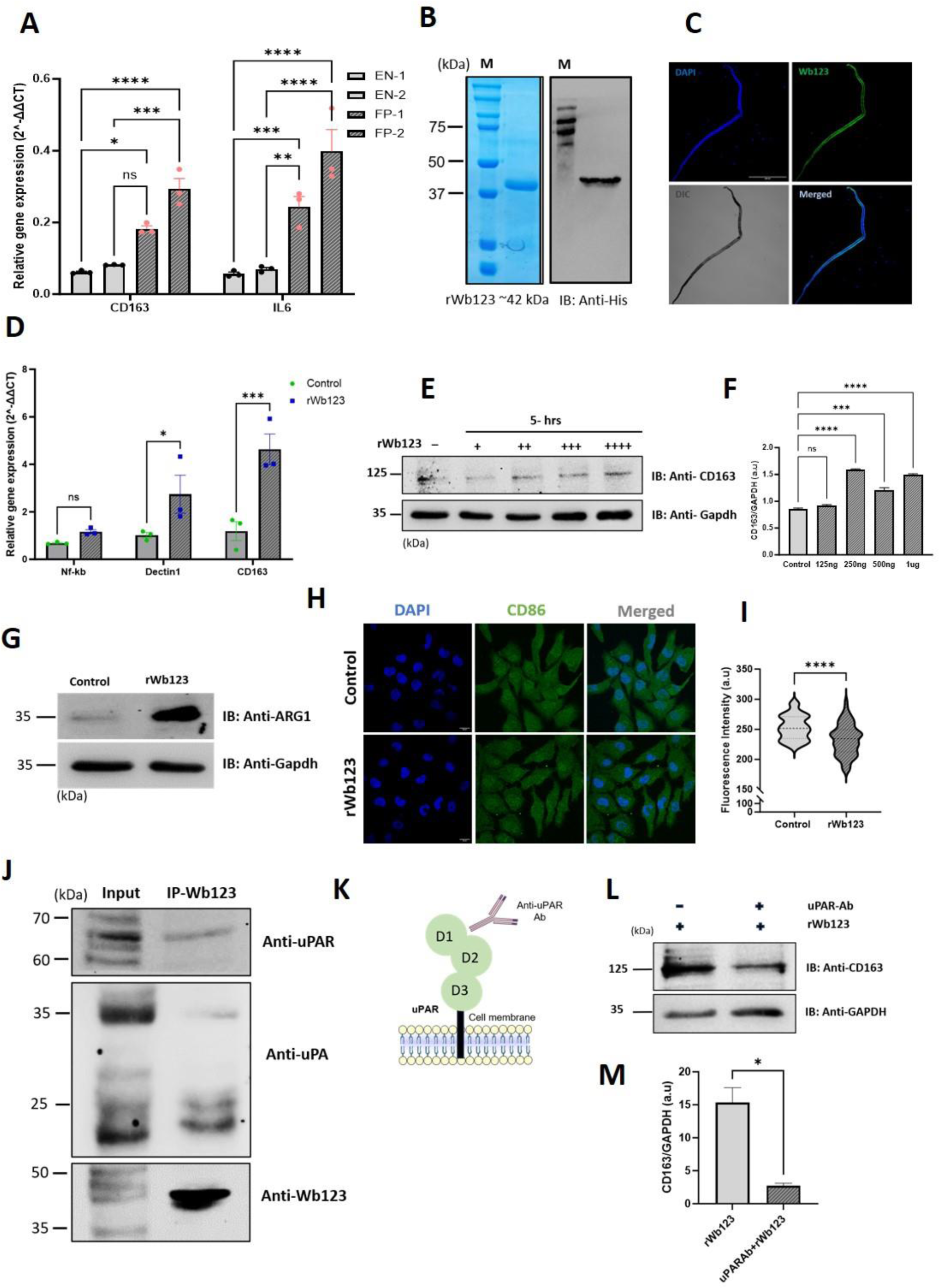
Wb123 induces alternate activation through uPAR. A) CD163 and IL6 expression in PBMCs of MF+ filarial patients. The graph represents fold change in gene expression of filarial patient compared with endemic normal. Data represent the mean ± SEM (n = 3). * Represents significance values compared to control, *p<0.0214, **p=0.0012, ***p=0.0006 and 0.0001, ****p<0.0001. Statistical significance was calculated using two-way ANOVA; B) SDS-PAGE stained with Coomassie brilliant blue dye and western blot analysis of purified rWb123 probed with His tag antibody; C) Immunofluorescence images showing Wb123 expression in microfilaria by confocal microscopy after incubation with monoclonal antibody; D) Study of activation marker expression after incubation with rWb123. The graph represents fold change in gene expression. Data represent the mean ± SEM (n = 3). * Represents significance values compared to control, *p<0.0194, ***p=0.0002 and 0.0005. Statistical significance was calculated using two-way ANOVA; E) Blots showing CD163 expression at protein level in rWb123 treated cells compared to control cells; F) Bar graph showing CD163 expression in different Wb123 concentrations represented as +(125ng), ++(250ng), +++(500ng), ++++(1µg). * Represents significance values compared to control, ***p=0.0008, ****p<0.0001. Statistical significance was calculated using two-way ANOVA; G) Blot showing arginase-1 expression in Wb123 treated cells and untreated cells; H) Immunofluorescence images showing CD86 expression in Wb123 treated cells and untreated cells; I) Intensity was determined from cells (Violin plot, n=70, * represents significance values compared to control, *P<0.0107, ****p<0.0001). Statistical significance was calculated using unpaired t test; J) Immunoprecipitation of Wb123 was performed from Wb123 treated HMC3 cells. Western blot analysis was performed with immunoprecipitated samples with uPA and uPAR as indicated in figure; K) Strategy for antibody blocking experiment. L) Blot showing CD163 expression in uPAR blocked cells and control cells; M) The bar graph represents quantification of CD163 expression in uPAR pre-treated and untreated cells. normalized to GAPDH (Mean± SEM, n=3, AU depicts arbitrary unit, * represents significance values compared to control, *p<0.00307).

Serpins not only acts as inhibitors of serine proteases but have also evolved alternative, non-inhibitory roles.^21^ Recombinant Wb123 (rWb123) was expressed in *Escherichia coli* as a 6x his tagged protein and purified by Nickel NTA affinity chromatography. The molecular weight of purified rWb123 was observed at 42kDa on SDS PAGE, confirmed through western blot analysis. (Fig. 2B) We also detected the expression of Wb123 in the outer membrane of microfilaria using an anti-Wb123 antibody. (Fig. 2C) To investigate the effect of rWb123 on macrophage activation, we performed qPCR analysis and found increased CD163 and Dectin-1 expression after rWb123 treatment compared to control cells. (Fig. 2D) Our western blot analysis confirmed increased CD163 expression after treatment with concentrations ranging from 125ng to 1µg. (Fig. 2E, F) Given that the induction of arginase-1 is crucial for establishing alternate activation, we investigated its expression levels and observed a significant increase in arginase-1 compared to control cells. (Fig. 2G, Supplementary Fig.1 C, D) Additionally, the classical activation marker CD86 was downregulated, confirming that Wb123 polarizes macrophage towards alternate activation. (Fig. 2H, I, Supplementary Fig. E) To identify the interacting partners of Wb123, we performed an immunoprecipitation assay using an anti-wb123 antibody followed by western blot analysis with Toll-like receptor 4 (TLR4), uPA and uPAR antibodies. (Fig. 2J) Our western blot results showed that Wb123 interacts with uPA and uPAR but not with TLR4. To determine whether Wb123 interaction with uPAR is necessary for its function, we performed an antibody-blocking experiment since uPAR is a GPI-anchored receptor. (Fig. 2K) We found that CD163 expression was significantly reduced in cells pretreated with uPAR antibody. (Fig. 2L, M) These results were further corroborated by immunofluorescence experiments, and no such effect on CD163 expression were observed in cells pre-incubated with LRP1 antibody. (Supplementary Fig. 1F, G)

### Wb123 mediated alternate activation is IL6 and STAT3 dependent

Cytokines such as IL-4, IL-10 and IL-13 are known to induce alternate activation of macrophages.^22,23^ We investigated which cytokine plays a major role in Wb123-induced alternate activation. Our qPCR results indicated that the expression of IL-6 and IL-8 was significantly higher in Wb123-treated cells compared with control cels. (Fig. 3A) Since STAT3 is a downstream signaling target of IL-6 and had been previously shown to play a major role in alternate activation.^20^ We analyzed and found that the expression of phosphorylated STAT3 was significantly higher in rWb123-incubated cells. (Fig. 3B, C) Additionally, we confirmed increased phosphorylation of STAT3 after incubation with concentrations ranging from 125ng to 1µg. (Fig. 3D) We then investigated the role of nuclear factor kappa-light-chain-enhancer of activated B cells (NF-kB), a known stimulator involved in IL6-mediated CD163 expression. We used an NF-kB cell-permeable inhibitory peptide (SN50i) and performed flow cytometry. Cells pre-treated with SN50i showed a 28-30 % reduction in CD163 expression. (Fig. 3E) Furthermore, we eliminated the possibility of LPS-induced NF-kB activation using TIRAP, a cell-permeable inhibitor. These results were further corroborated by western blot analysis. (Supplementary Fig. 1H)

**Figure 3:**
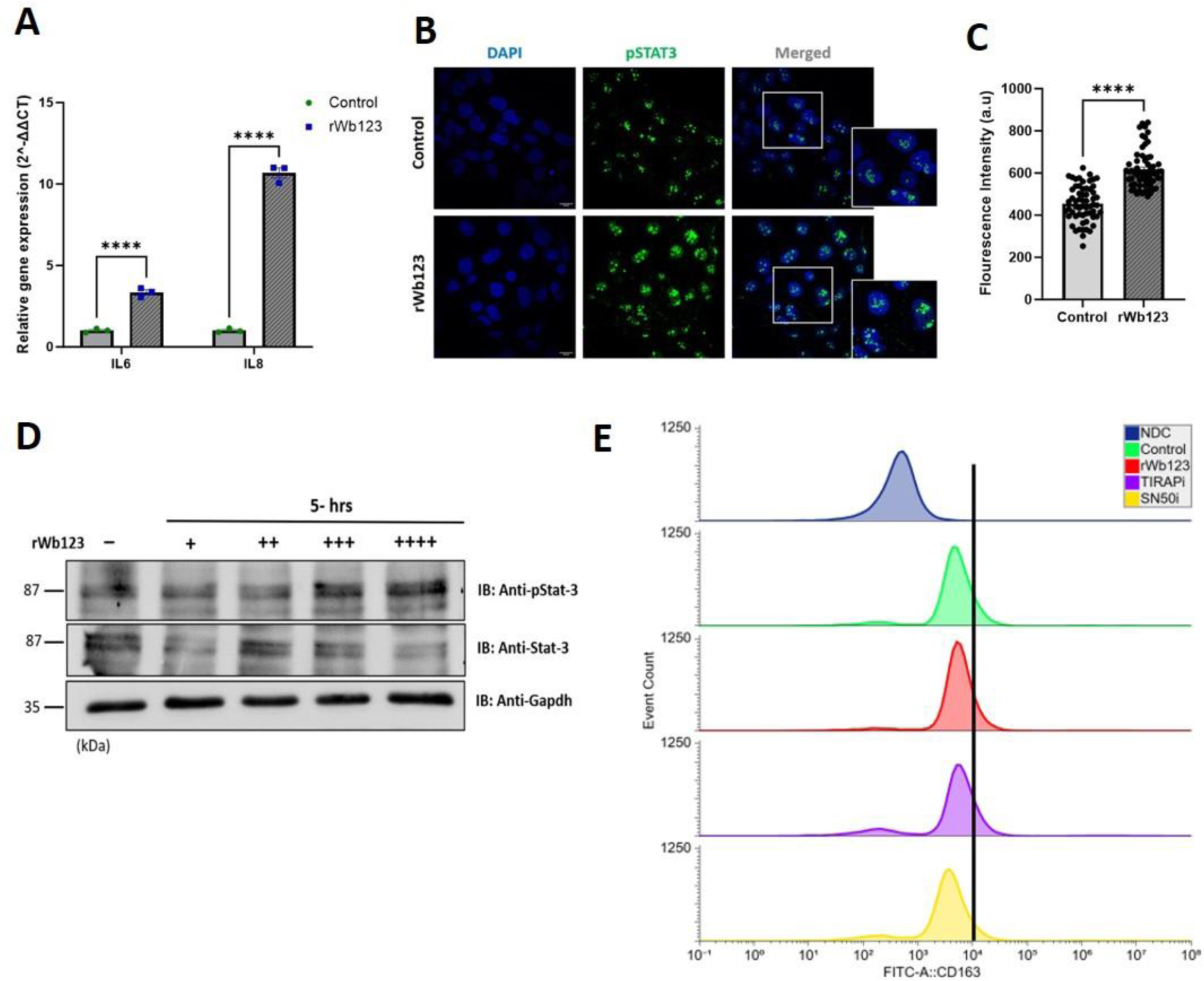
Wb123 induced alternate activation is IL-6-STAT3 mediated. A) Study of cytokine expression after incubation with rWb123 protein. The graph represents fold change in gene expression. Data represent the mean ± SEM (n = 3). * Represents significance values compared to control, ****p<0.0001. Statistical significance was calculated using two-way ANOVA; B) Immunofluorescence images showing phosphorylated STAT3 expression in Wb123 treated and untreated cells; C) Intensity was determined from cells (Mean± SEM, n=60, ****p<0.0001). Statistical significance was calculated using unpaired t test; D) Blots showing concentration dependent increase in phosphorylated STAT3 expression at protein level in rWb123 treated cells compared to control cells; E) Stacked histogram representing CD163 expression in control cells, rWb123 treated cells and cells treated with TLR4 peptide inhibitor TIRAPi and NF-kB peptide inhibitor SN50i respectively.

### Wb123 dysregulates the LPS-IFN-γ induced classical activation and oxidative stress

As discussed in previous results, Wb123 induces alternate macrophage activation suggesting that it could dysregulate classical activation of macrophage. To evaluate the effect of Wb123 on classical activation, we pre-incubated cells with LPS and IFN-γ, collectively referred to as LPS-I, well-known inducers of classical activation.^24^ This was followed by rWb123 treatment. (Fig. 4A)

**Figure 4:**
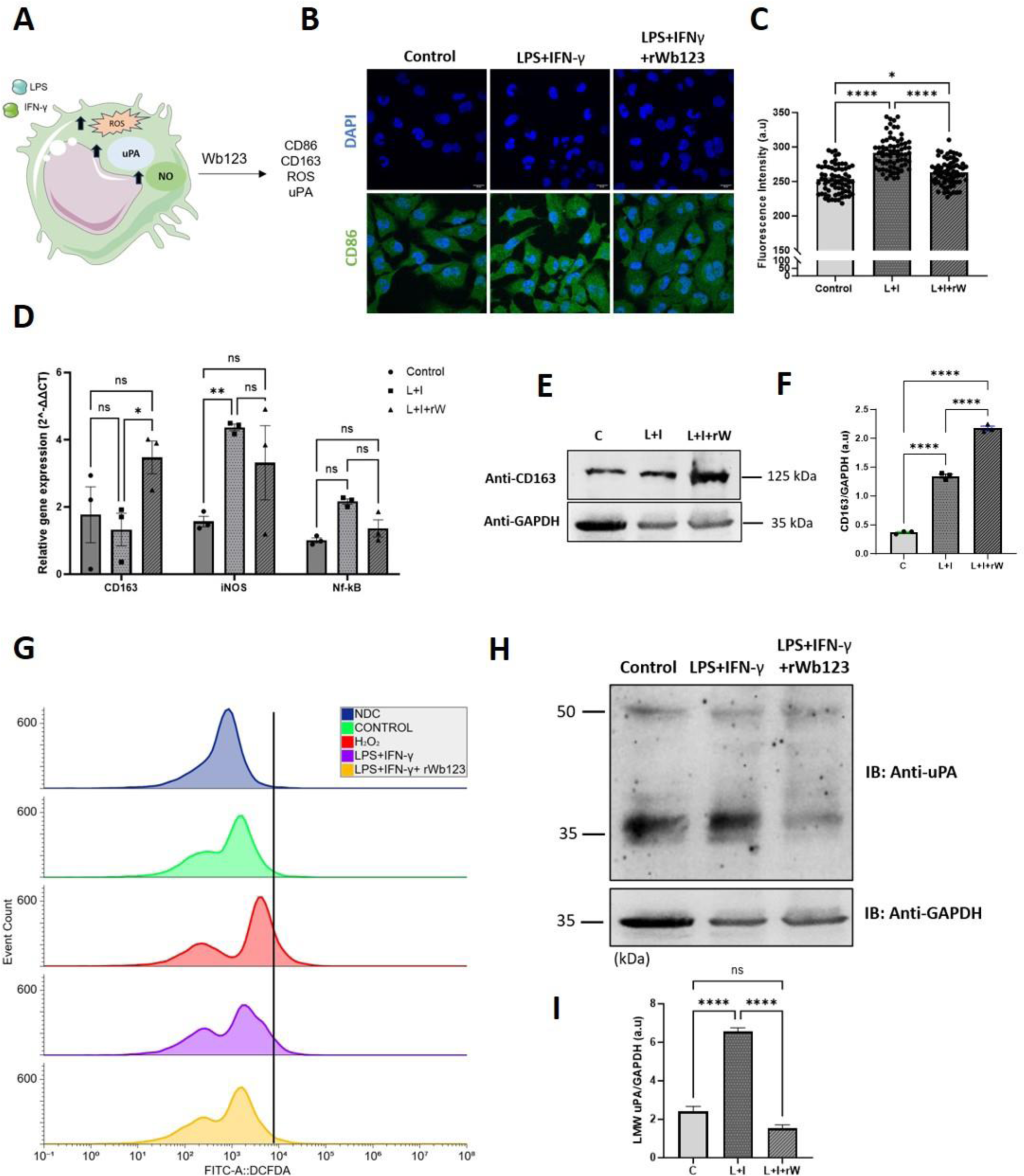
LPS-I induced classical activation is abrogated by Wb123. A) Experimental strategy for evaluating Wb123 effect on LPS-I induced pro-inflammatory response. B) Immunofluorescence images showing CD86 expression in control cells compared with LPS-I and LPS-I treated cells with Wb123; C) Intensity was determined from cells (Mean± SEM, n=70, * represents significance values compared to control, *P<0.0132, ****p<0.0001). Statistical significance was calculated using one-way ANOVA; D) Activation marker expression after incubation with LPS-I and LPS-I with Wb123. The graph represents fold change in gene expression. Data represent the mean ± SEM (n = 3). * Represents significance values compared to control, ****p<0.0001. Statistical significance was calculated using two-way ANOVA; E) Blots showing CD163 expression in control cells compared with LPS-I and LPS-I with Wb123; F) The bar graph represents quantification of CD163 expression normalized to GAPDH (Mean± SEM, n=3, AU depicts arbitrary unit, * represents significance values compared to control, ****p<0.0001); G) Stacked histogram representing ROS positive cells in control cells compared to LPS-I and LPS-I with Wb123, H2O2 as a positive control; H) Blots showing uPA expression in control cells compared with LPS-I and LPS-I with Wb123; I) The bar graph represents quantification of LMW uPA (∼30kDA) expression normalized to GAPDH (Mean± SEM, n=3, AU depicts arbitrary unit, * represents significance values compared to control, ****p<0.0001);

As expected, our qPCR results indicated that LPS-I treated cells showed no effect on CD163 expression, however, CD163 expression was significantly higher in the cells treated with both LPS-I and Wb123. (Fig. 4B, C) Furthermore, we validated the increased CD163 and decreased CD86 expression using immunofluorescence and western blot studies. (Fig. 4E, F, Supplementary Fig. 2A, B) Given that Wb123 may dysregulate the oxidative status of the cells, we probed ROS production by labelling cellular ROS with H_2_DCFDA. Flow cytometry results revealed a 15-17% increase in ROS production after LPS-I treatment which decreased to 7-9% following rWb123 incubation. Additionally, cells treated with 20µm hydrogen peroxide (H_2_O_2_) showed a 32-34 % increase in ROS production. (Fig. 4G, Supplementary Fig. C, D) We also evaluated the effect on NO production and found that NO production was significantly decreased in cells incubated with rWb123, even after LPS-I activation.^25^ (Supplementary Fig. 2E, F) uPA expression has been correlated with the oxidative status of macrophages and is known to induce ROS production.^26^ Therefore, we explored the effect of Wb123 on uPA expression and found that low molecular weight uPA expression was significantly reduced in rWb123-incubated cells, even after LPS-I activation. (Fig. 4H, I) We further validated these results through immunofluorescence and western blot studies. (Supplementary Fig. 2G, H)

### Anti-Wb123 antibody abrogates the effects of Wb123 on macrophage activation

We used the anti-wb123 monoclonal antibody ‘MabG8’ to investigate its effect on Wb123 through an antibody neutralization assay. (Fig. 5A, Supplementary Fig. 3A, B) Our results indicated that overnight incubation of MabG8 with rWb123 reduces Wb123-induced CD163 expression, with rWb123-treated cells serving as a control. (Fig. 5B, Supplementary Fig. 3C, D) Furthermore, we examined the effect of the monoclonal antibody on LPS-I-induced ROS production by labeling cellular ROS with H_2_DCFDA. As expected, our results suggested that Wb123 decreases the ROS levels. (Fig. 5C, D) However, MabG8-Wb123 did not reduce the ROS levels. Additionally, uPA expression was unaffected by the MabG8-Wb123 treatment, while it was decreased by rWb123 treatment. (Fig. 5E, F) Overall, our data illustrates that the monoclonal antibody MabG8 hinders the anti-inflammatory effects of Wb123 and reduces CD163 expression.

**Figure 5:**
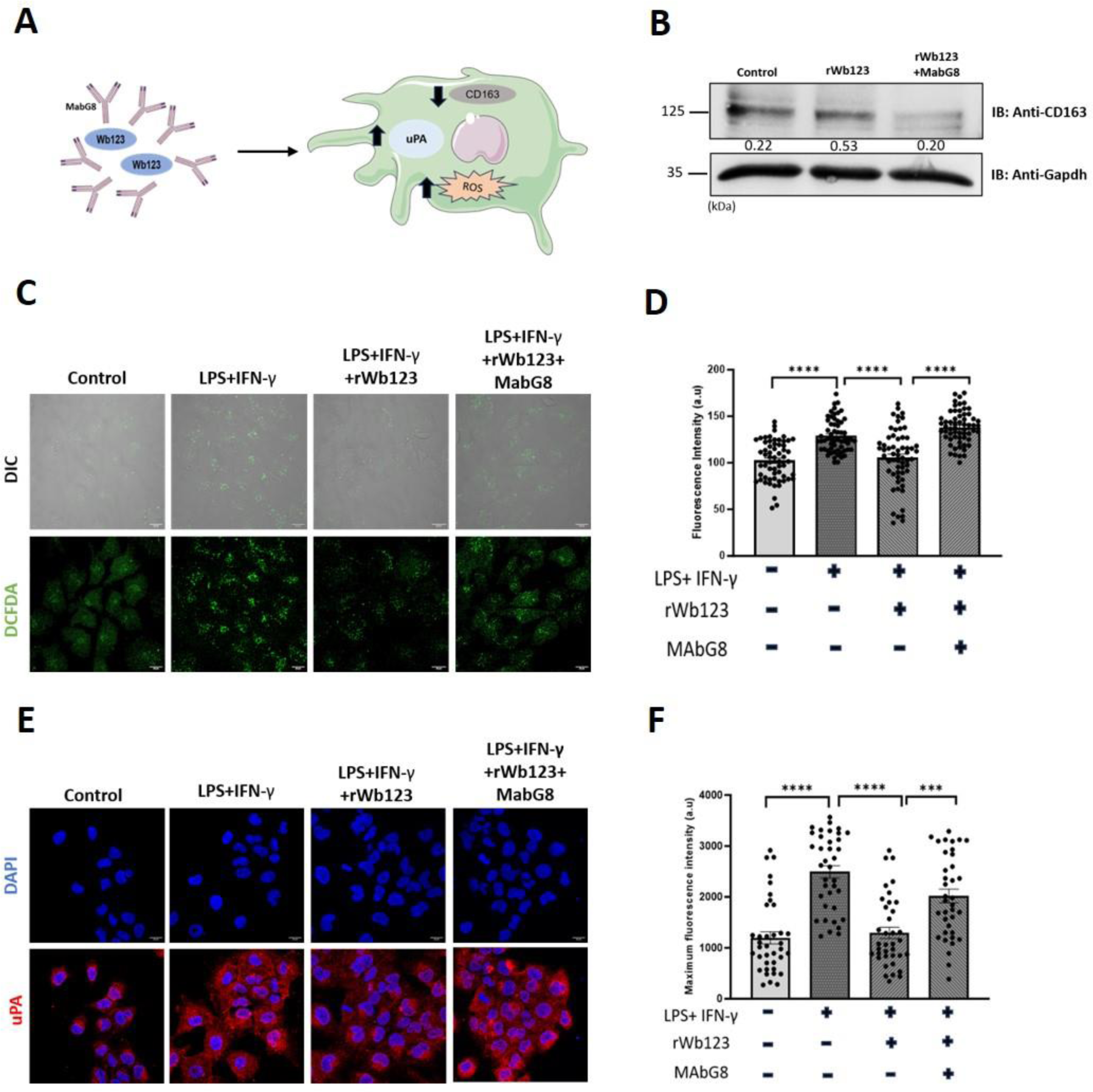
Monoclonal antibody MAbG8 impedes the Wb123-induced alternative activation. A) Experimental strategy to evaluate effects of monoclonal antibody; B) Blots showing CD163 expression at protein level in control cells compared with cells treated with rWb123 and rWb123 with monoclonal antibody (MAbG8); C) Confocal microscopy images showing ROS expression in control cells compared with LPS-I, LPS-I with rWb123 and LPS-I with rWb123-MAbG8; D) Intensity was determined from cells (Mean± SEM, n=80, ***p=0.0003, ****p<0.0001); E) Immunofluorescence images showing uPA expression in control cells compared with LPS-I, LPS-I with rWb123 and LPS-I with rWb123-MAbG8; F) Intensity was determined from cells (Mean± SEM, n=60, ns=0.1120, ****p<0.0001). Statistical significance was calculated using ordinary one-way ANOVA.

## 4. Discussion

Filarial worms possess a remarkable ability to modulate the host immune response, characterized by downregulated T cell response and muted macrophage response. While the reason for downregulated T-cell response has been frequently investigated, the understanding of macrophage activation remains scarce. Using publicly available data, we demonstrated that filarial patients exhibit increased expression of alternative activation markers and decreased expression of classical activation markers in peripheral monocytes, with these changes reversing after treatment.(Supplementary Fig.).^12^ A recent report highlighted that the serine protease inhibitor Ts-serpin plays an essential role in macrophage alternate activation.^17^ Using in-silico analysis, we identified fifteen putative filarial serpins with conserved RCL including Wb123 which is expressed at microfilaria stage. We also validated that serpin Wb123 induces the alternate activation of macrophages, as shown by increased CD163, Dectin-1 and Arginase-1 expression. Additionally, we observed that Wb123 induces the IL6 and Il8 expression, cytokines not generally associated with alternate activation. From our literature review, we found that a human serpin protein (SERPINE1) induces IL6 upregulation and subsequent activation of STAT3 through p38 MAPK and NF-kB pathways.^20^ Our results validated that Wb123 increases the STAT3 phosphorylation, which in turn induces CD163 expression. This process is dependent on NF-kB and occurs independently of TLR4-mediated NF-kB activation. Notably, several reports have shown that inducers like LPS-I induces a robust classical activation state marked by increased expression of pro-inflammatory markers.^8,24^ Our study indicated that Wb123 dysregulates the LPS-I-induced classical responses, as demonstrated by a decrease in CD86 expression and an increase in CD163 expression. We also found that Wb123 downregulates ROS and NO production which are essential mediators in pathogen elimination and promoting inflammation. Furthermore, we validated that LPS-I incubation upregulates the uPA expression as a part of the immune response. However, our study indicated that Wb123 downregulates the uPA expression, specifically the catalytically active low molecular uPA(35kDa), which is reported to cleave uPA receptor to produce soluble uPAR a known marker of chronic inflammation.^27,28^ Thus, Wb123 not only induces alternate activation of macrophages but also affects the pro-oxidative function of macrophages essential for pathogen elimination. We evaluated the effect of monoclonal antibody-based neutralization on Wb123 function and our results suggested that anti-Wb123 monoclonal antibody decreases the CD163 expression while rescuing uPA and ROS expression after monoclonal antibody incubation.

Interestingly, an immunoprecipitation assay using the anti-Wb123 antibody suggested that Wb123 interacts with uPA and uPAR. However, the exact mechanism by which wb123 initiates the alternate activation of macrophage remains unknown. Additionally, our results suggested that uPAR plays a major role in Wb123-induced alternate activation since antibody-based blocking of uPAR reduced CD163 expression. However, uPAR is a GPI linked receptor that cannot induce signaling unless it interacts with another transmembrane receptor within uPAR interactome such as EGFR and integrins requiring further exploration.^29^ A study of *Brugia malayi* L3 larva infection in mice showed alternate activation of macrophage as early as day 3 post-infection while IL10 expression was not detected until day 7 post infection.^30^ Filarial serpin-induced alternate activation might play significant role in early alternate macrophage activation. Previous studies on filarial serpins have focused on exploring their inhibitory role on host serine proteases. One such study identified *Brugia malayi* microfilaria serpin BmSpn-2 as capable of modulating immune response by inhibiting human neutrophil elastase.^31^ However, subsequent research revealed their inability to form a stable complex with serine proteases.^32^ Although our fluorescence based inhibition assay indicated that Wb123 might inhibit catalytic activity of uPA. however, further investigation is necessary to assess its capability of form stable serpin-protease complexes. (Supplementary Fig. 3E) In conclusion, our results provide compelling evidence that filarial serpins exhibit significantly higher expression levels in L3 larval and microfilariae stages. We also identified and characterized a novel serine protease inhibitor from *Wuchereria bancrofti* (Wb123), which induces alternative activation of macrophages through a uPAR-dependent IL-6/STAT3 pathway while downregulating classical activation induced by LPS-I. Additionally, we demonstrated that the monoclonal antibody MAbG8 impedes the effects of Wb123, underscoring the potential of monoclonal antibody therapy for filarial patients. This study offers deeper insights into how filarial serpins modulate macrophage activation during the early stages of infection.

### Statistical analysis

All data points were collected for the calculation of standard error mean (SEM). Statistically significances were calculated based on experimental parameters by using unpaired t-test, one-way ANOVA and two-way ANOVA. P-Values and statistical test were mentioned on the respective figure.

## Acknowledgment

Ph.D. student P.U is grateful to the University Grant Commission (UGC) for fellowship and Ph.D. students AM and SM are grateful to Shiv Nadar Institute of Eminence Deemed to be University (SNIoE), Delhi NCR, and the Shiv Nadar Foundation for their research fellowships. The authors acknowledge the DST-FIST grant [SR/FST/LS-1/2017/59(c)] for the confocal microscopy facility at Shiv Nadar Institute of Eminence Deemed to be University, Delhi NCR and Shiv Nadar Core Research Grant.

## Authors contribution

Conceptualization: PU, SP and SS; Investigation and methodology: PU, AM, SM, AM, GK, MK, PG; Data analysis and Manuscript writing: PU, Manuscript Editing: SS, SP; Funding acquisition and supervision: SS, SP; All authors have read and agreed to the final version of the manuscript.

## Data availability statement

Data will be available on request

