## Supplementary Material for "Monoclonal antibody to Filarial serpin Wb123 impedes the urokinase plasminogen activator receptor mediated Alternative activation of macrophages"

**Supplementary figures-**

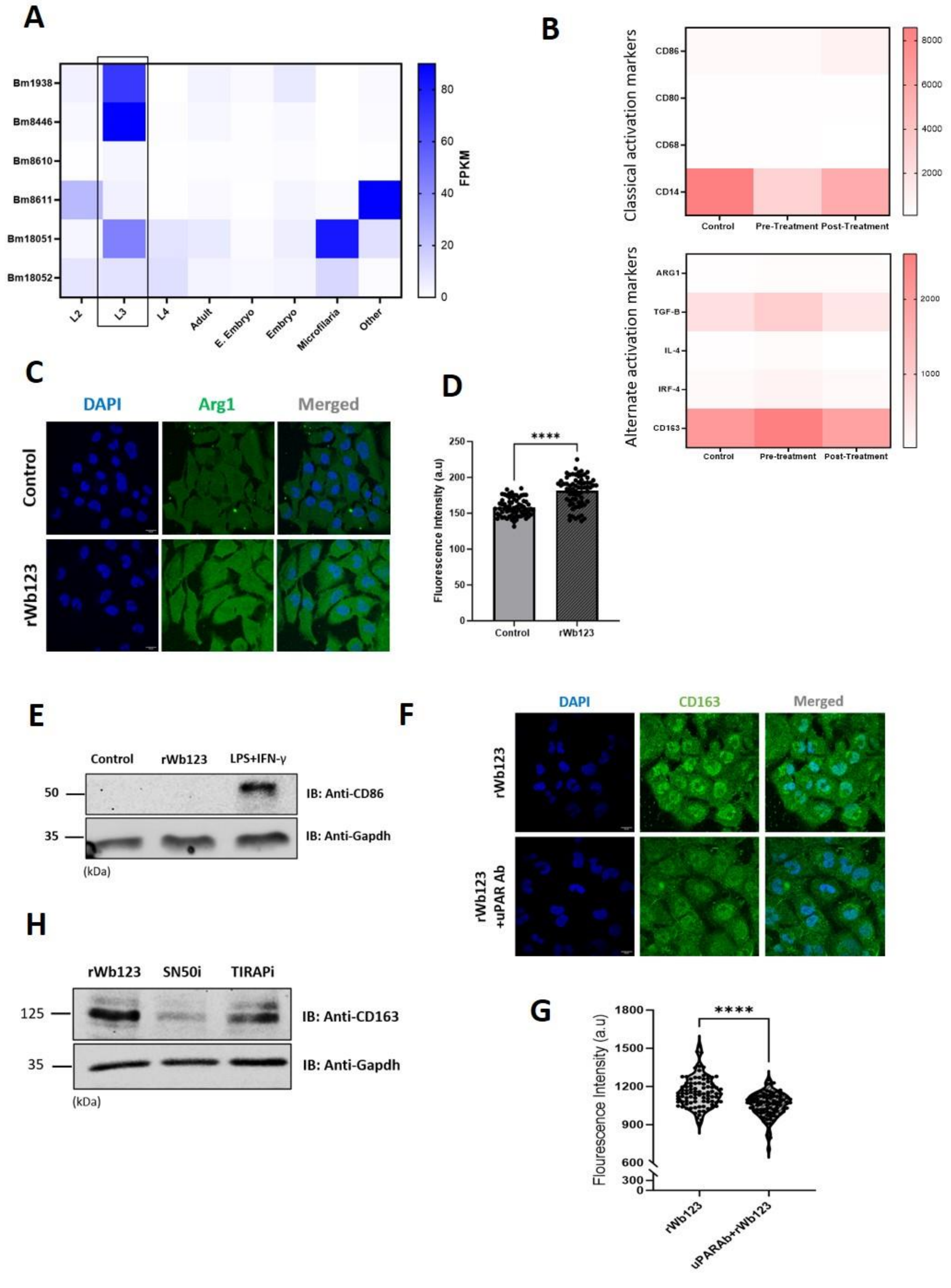

**Supplementary figure 1:**

A) Heatmap represents the expression of identified *Brugia malayi* serpins in different life stages from WormBase expression database; B) Heatmap represents expression of alternate and classical activation markers in pre-treatment, post-treatment and healthy person monocytes; C) Immunofluorescence images showing Arg-1 expression in Wb123 treated cells and untreated cells; D) Intensity was determined from cells (mean ± SEM, n=70, * represents significance values compared to control, ****$p<0.0001$). Statistical significance was calculated using unpaired t test; E) Blots showing CD86 expression in control cells compared with Wb123 and LPS-IFN-γ; F) Immunofluorescence images showing CD163 expression after Wb123 incubation in uPAR Ab pre-treated and untreated cells; G) Intensity was determined from cells (Violin plot, n=90, * represents significance values compared to control, ****$p<0.0001$). Statistical significance was calculated using unpaired t test; H) Blots showing CD163 expression in Wb123 treated cells compared with cells pre-treated with NF-kB and TLR4 peptide inhibitors.

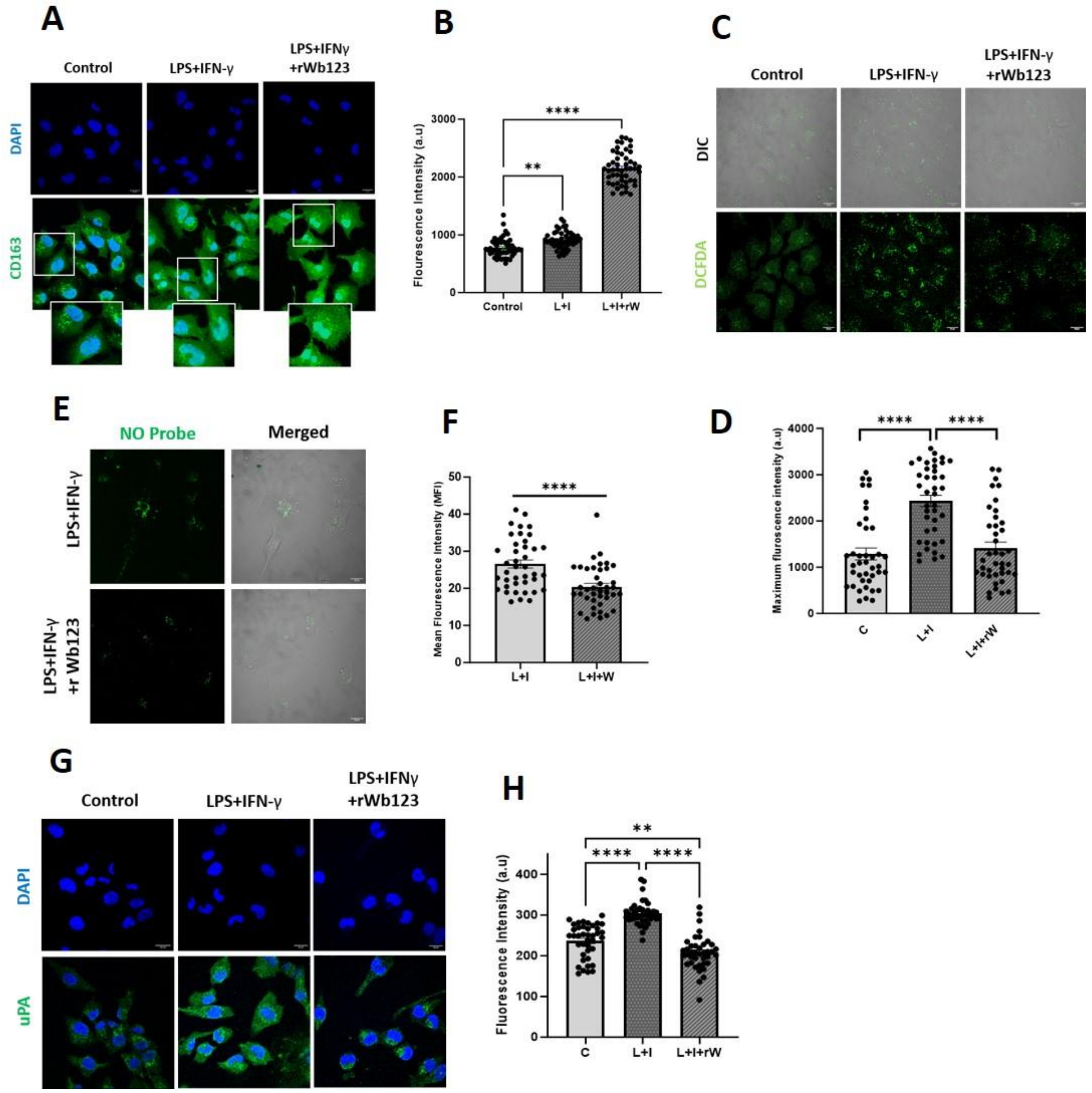


**Supplementary figure 2:**

A) Immunofluorescence images showing CD163 expression in control cells compared with cells treated with LPS-IFN-γ and LPS-IFN-γ with Wb123; (ii) Intensity was determined from cells (Mean± SEM, n=70, *p=0.0132, ****p<0.0001), Statistical significance was calculated using ordinary one-way ANOVA; C) Confocal microscopy images showing ROS expression in control cells compared with LPS-IFN-γ, LPS-IFN-γ with rWb123; (D) Intensity was determined from cells (Mean± SEM, n=40, ****p<0.0001); E) Images showing NO expression in LPS-IFN-γ treated cells compared to LPS-IFN-γ with rWb123 treated cells; F) Intensity was determined from cells (Mean± SEM, n=40, ****p<0.0001); G) Immunofluorescence images showing uPA expression in control cells compared with LPS-

IFN-γ and LPS-IFN-γ with Wb123; H) Intensity was determined from cells (Mean± SEM, n=40, **p=0.0025, ****p<0.0001); Statistical significance was calculated using ordinary one-way ANOVA;

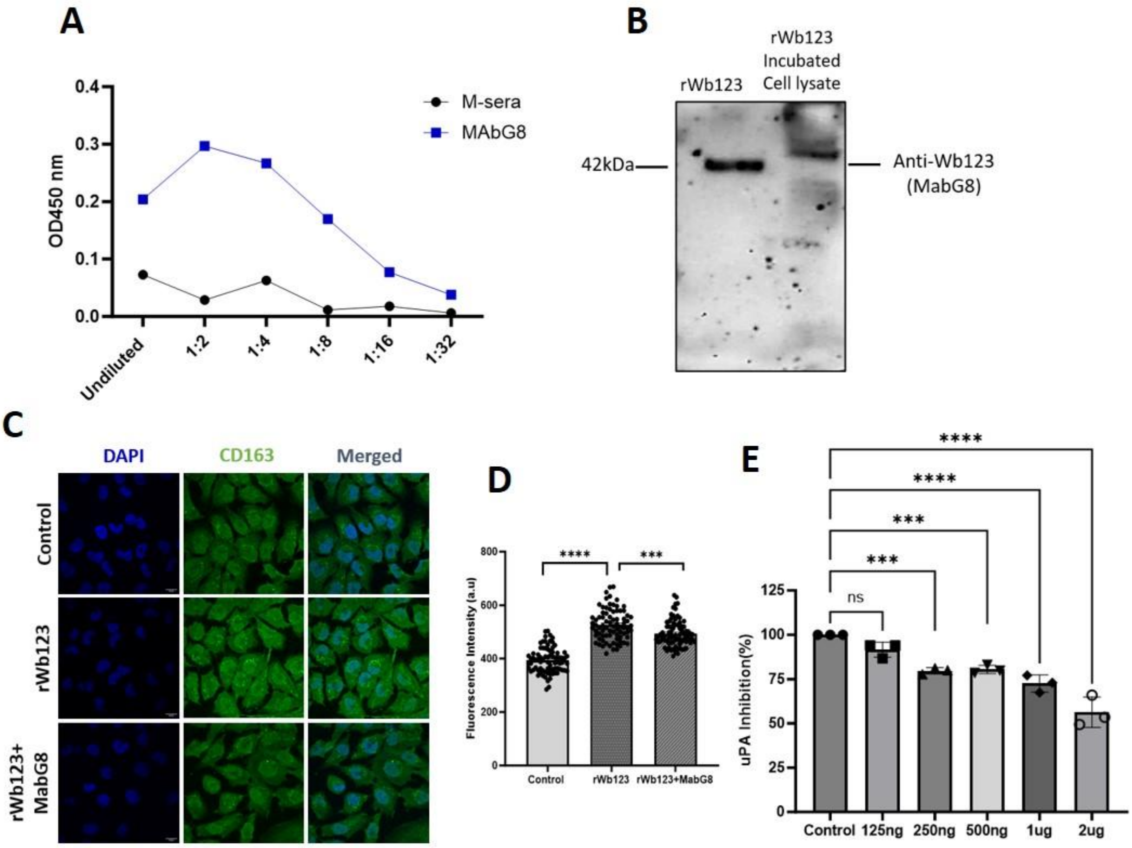


**Supplementary figure 3:**

A) Graph showing anti-Wb123 monoclonal antibody MabG8 binding with recombinant Wb123 using ELISA; B) Blots shows Wb123 protein in cell lysate 5 hours after incubation using MAbG8, rWb123 taken as control; C) Immunofluorescence images showing CD163 expression in control cells compared with Wb123, Wb123 with MAbG8; D) Intensity was determined from cells (Mean± SEM, n=80, ***p=0003, ****p<0.0001). Statistical significance was calculated using ordinary one-way ANOVA; E) Bar graph showing concentration dependent inhibition of uPA catalytic activity using recombinant uPA and fluorescence substrate;

**Table S1.** Wb123 structure quality check using Saves server and ProSa web.

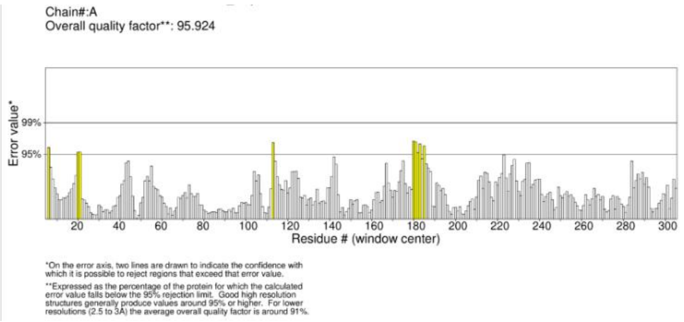


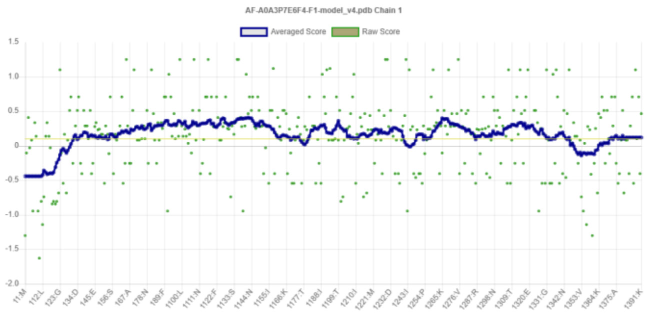


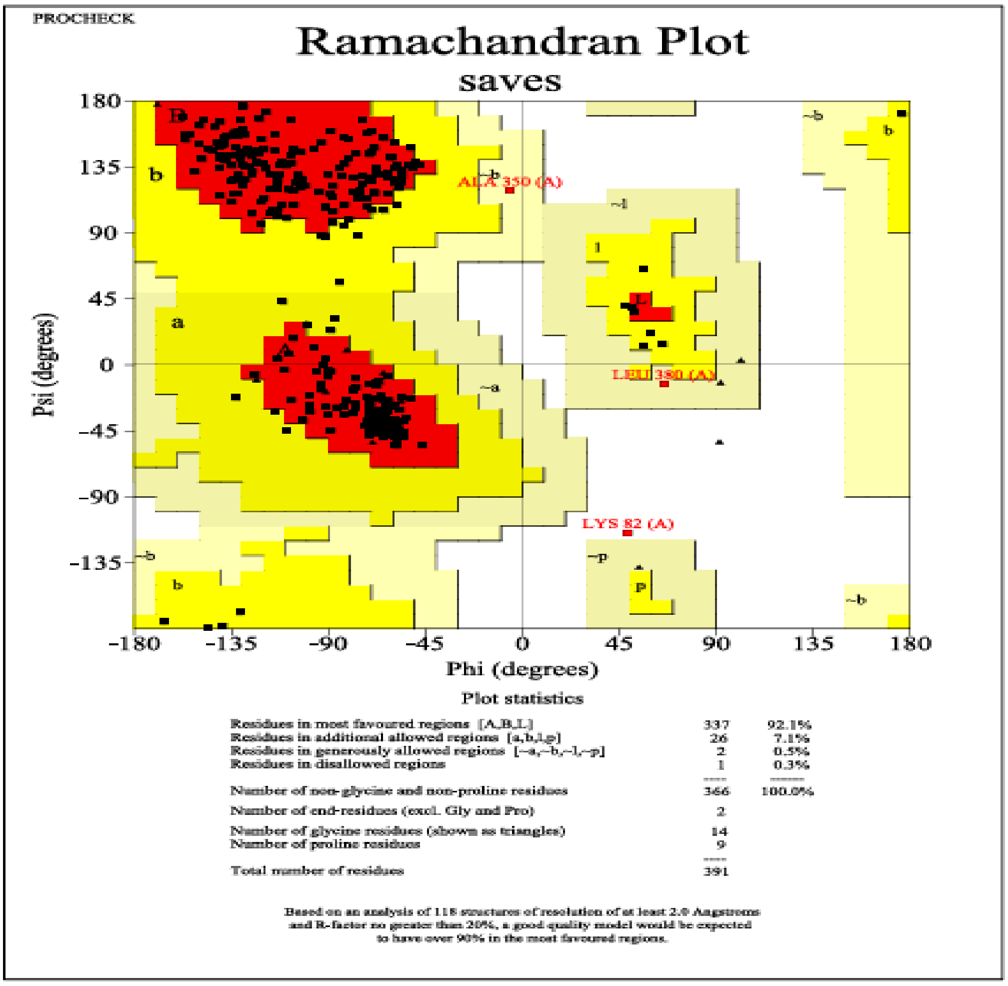


**Table S2.** Superimposed structure of Wb123 (Green) with resolved structure of *Homo sapiens* serpins (Cyan).

| | **Resolved Serpin Structures** | **PDB ID** | **RMSD (Å)** | **Superimposed Structure** |
|---|---|---|---|---|
| | Plasminogen Activator Inhibitor-1 | 3PB1 | 1.530 | 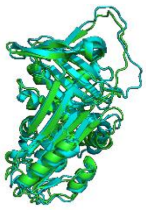 |
| | Alpha-1-Antitrypsin | 1HP7 | 1.687 | 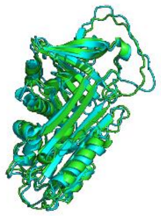 |
| Wb123 (Wuchereria bancrofti) | Kallistatin | 6F02 | 1.845 | 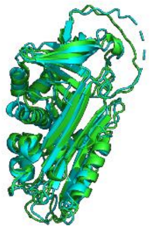 |
| | MASPIN | 1XQP | 1.841 | 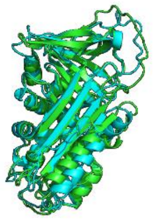 |
| | Glia derived nexin | 4DY0 | 1.611 | 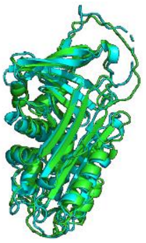 |
| | Serpin B3 | 2ZV6 | 1.834 | 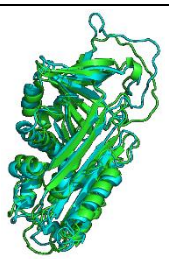 |
